# FORGE: A tool to discover cell specific enrichments of GWAS associated SNPs in regulatory regions

**DOI:** 10.1101/013045

**Authors:** Ian Dunham, Eugene Kulesha, Valentina Iotchkova, Sandro Morganella, Ewan Birney

## Abstract

Genome wide association studies provide an unbiased discovery mechanism for numerous human diseases. However, a frustration in the analysis of GWAS is that the majority of variants discovered do not directly alter protein-coding genes. We have developed a simple analysis approach that detects the tissue-specific regulatory component of a set of GWAS SNPs by identifying enrichment of overlap with DNase I hotspots from diverse tissue samples. Functional element Overlap analysis of the Results of GWAS Experiments (FORGE) is available as a web tool and as standalone software and provides tabular and graphical summaries of the enrichments. Conducting FORGE analysis on SNP sets for 260 phenotypes available from the GWAS catalogue reveals numerous overlap enrichments with tissue–specific components reflecting the known aetiology of the phenotypes as well as revealing other unforeseen tissue involvements that may lead to mechanistic insights for disease.

## Identifying Regulatory Components of Genome Wide Association Study Hit Lists

A primary motivation for sequencing the human genome was to shed light on mechanisms involved in human disease. Since finishing the sequence there has been much activity in two areas towards that goal. In the first, extensive re-sequencing of individual genomes has provided comprehensive lists of human variations, which can in turn be examined for association with disease and other phenotypes in Genome Wide Association Studies (GWAS) [1]. In the second area, efforts have been undertaken to identify the specific sequences that enact function within the genome including, but not restricted to, regions defining genes and their controlling elements [2–4]. The aim, of course, is to understand the associations uncovered in the first approach in the context of the annotations delivered from the second.

The past few years have seen a dramatic growth in the number of variants associated with disease by GWAS [1]. An extensive catalogue of GWAS associations has been compiled containing nearly 14,000 associations of variants to phenotypes [5]. However, a crucial observation is that the majority of the variants observed do not directly affect the coding regions of protein coding genes. Notwithstanding that the reported variant for an association may be in linkage disequilibrium with a causal variant affecting a protein coding sequence, regulatory regions have been demonstrated to be linked to both specific diseases associations [6–18] (see [19] for review and further examples) and to be enriched in bulk in SNPs found across all GWAS [2], [20–22]. The ENCODE consortium reported that GWAS single nucleotide variants are substantially enriched in regulatory regions and up to 80% of GWAS variants have a potential regulatory interpretation via overlap with regulatory annotation [2], [21], [22]. Furthermore, Maurano et al [21] showed that regulatory regions revealed by the DNase-seq method show a cell specific enrichment for GWAS variants in specific phenotypes consistent with probable physiological mechanisms. Trynka et al [23] similarly found that regulatory elements identified by the histone modification H3K4me3 show a phenotypically relevant cell specific overlap with GWAS SNPs. Several tools exist to highlight the specific overlaps of individual GWAS SNPs with potential regulatory regions [24], [25]. To date however, much of the focus on this work has been on prioritising variants, rather than exploring the extensive cell type information present in the large-scale projects.

We have developed a simple but powerful approach that identifies significant cell specific enrichments in regulatory regions for sets of single nucleotide variants, typically from GWAS. We name the approach Functional element Overlap analysis of the Results of GWAS Experiments or FORGE, and have implemented it as both a rapid web tool for ENCODE[26] and Roadmap Epigenome project DNase-seq data[27] and a free-standing open source software. The web tool produces two alternative graphical outputs for exploration alongside tabulated enrichment data. FORGE analysis across all eligible phenotypes in the entire GWAS catalog [5] identifies numerous interesting patterns of enrichment by cell type and suggests tissues to focus on for future follow up studies.

## Forge Analysis approach

FORGE analysis provides a method to view the tissue specific regulatory component of a set of variants. In its current implementation, FORGE analysis takes a set of single nucleotide polymorphisms (SNPs), such as those SNPs reported above the genome wide significance threshold (p < 5e-8) in a GWAS study, optionally filters the SNPs to remove all bar one SNP from a region in high LD (“LD pruning”) and determines whether there is enrichment for overlap with putative regulatory elements compared to a matched background of SNP sets. Initially the elements considered are DNase I hotspots generated from either the ENCODE [26] or Roadmap Epigenomics projects DNase I data by the Hotspot method [28, 29], because of both the comprehensiveness of the sites identified and the broad range of cell types for which DNase I data was available. DNase I hotspots can be regarded as regions of general DNase I sensitivity.

For each set of test SNPs, an overlap analysis is performed against the DNase I hotspots for each available cell sample separately (125 samples for ENCODE, 299 for Roadmap, described in Supplementary Table S1), and the number of overlaps is counted. Major potential confounders in this analysis are the many biases of GWAS SNP distribution on the genome. To account for this a background distribution of the expected overlap counts for this SNP set is obtained by identifying 1000 matched background SNP sets of the same number of SNPs, matching each test SNP with an equivalent SNP by decile bin for each of G+C content (GC), minor allele frequency (maf) and distance to the nearest transcription start site (TSS). The matched background SNPs sets are overlapped with the DNase I hotspots and the background distribution of overlap counts is determined. The enrichment of the test SNP set for each sample is expressed as the binomial P value of the test SNP set given the background overlap distribution. The FORGE results are presented in interactive and static graphical and tabular forms by cell type. Enrichments above the background distribution with binomial P values less than 0.01 corrected for multiple testing are considered significant and are highlighted in red in the graphical output. Enrichments with p <= 0.05 are also highlighted in pink (Figure 1). As the DNase I patterns are not independent between cell types, we conducted simulation experiments with randomly selected input SNPs. We chose 1000 random test SNP sets for each of a series of SNP counts ranging between 5 and 100 SNPs and conducted FORGE analysis on both ENCODE and Roadmap data. The false positive rate was determined as the number of cell type enrichments identified greater than the significance thresholds used by FORGE expressed as the proportion of the total number of sample overlap tests performed (424,000) for each SNP count. This analysis showed that the P value thresholds are reasonably well calibrated to false positive levels around 0.5-0.75 %. In practice we correct the thresholds further for multiple testing across different tissues so as to be more stringent and so expect typical false positive levels to be less. As discussed below, many of the GWAS SNP sets do not reveal any enrichment, consistent with low false positive rates.

## The FORGE tool

We have implemented FORGE as a web tool available at http://browser.1000genomes.org/Homo_sapiens/UserData/Forge. The interface accepts a list of SNPs by dbSNP RefSNP identifier (RSID) or by genomic location on human genome build GRCh37 in either Variant Call Format (VCF, http://www.1000genomes.org/wiki/Analysis/Variant%20Call%20Format/vcf-variant-call-format-version-41) or Bed format (Personal Genome SNP format, http://genome.ucsc.edu/FAQ/FAQformat.html#format10), and allows specification of the background selection from two common sets of GWAS SNP typing microarrays. LD filtering is achieved at either r^2^ >= 0.8 or r^2^ >= 0.1 using 1000 genomes project population data. The outputs of the analysis are an interactive graphic for exploration of the analysis, a static pdf for printing or publication (Figure 1), and a table of enrichments in either an interactive or standard tab separated format.

In addition the code is available to download from https://github.com/iandunham/Forge, with the required database files available at ftp://ftp.1000genomes.ebi.ac.uk/vol1/ftp/technical/browser/forge. Installed on a Macbook Pro with core i7 processor, 16Gb RAM and a solid state hard drive, a typical FORGE analysis of a set of 55 SNPs with 1000 background tests is accomplished in around 30s, and in 35s with LD filtering.

## Gallery of Examples

We ran FORGE analysis on 260 phenotypes in the NCBI GWAS catalog NCBI GWAS catalog [5, 30] with a reported associated SNP count of 5 or more after LD pruning (see Supplementary Table S2 for list of phenotypes analysed and references) at genome-wide significance. Complete tables of results for all SNP sets analysed are included in the data directory of the Github release, https://github.com/iandunham/Forge. 35 and 60 out of 260 SNP sets had at least one significant enrichment at the P value thresholds of 0.05 and 0.01 after correction for multiple testing, respectively (Table 1). A set of example positive outputs from this analysis is available from http://www.1000genomes.org/forge-gwas-catalog-example-gallery11 (PDF format). Removing SNPs that directly alter a protein coding exon from the GWAS catalog sets did not substantively alter the patterns of enrichments (data not shown).

**Table 1:**
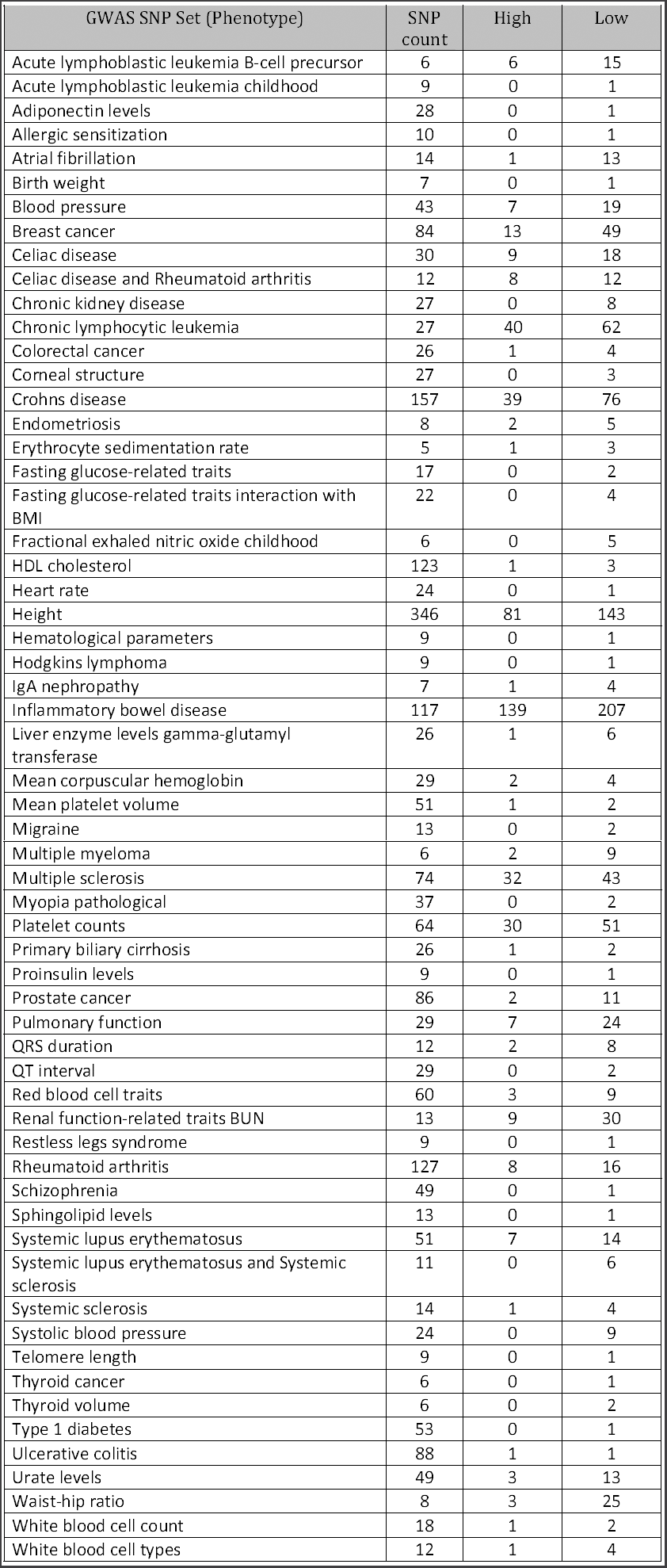
A list of SNP sets with positive enrichments. GWAS SNP set gives the phenotype of the study for which these SNPs were found to be associated as recorded in the GWAS catalog. SNP Count is the number of SNPs analysed before LD pruning. High and Low columns give the number of DNas1 cell samples found to be enriched for overlap in the Forge analysis at binomial p >= 0.01 (High) and p >= 0.05 (Low) thresholds. Further details of the SNP sets analysed are given in Supplementary Table S2.

Figure 1 shows a series of example FORGE analyses for autoimmune disease studies on the Roadmap Epigenome samples (references for the studies are provided in Supplementary Table S2). In each case there is a clear signal for enrichment of overlap with DNase I hotspots in the blood-derived samples including cells of immune function. In more detail, for those phenotypes where there is involvement of T cell activation or invasion in the aetiology (e.g. Crohn’s disease, Multiple Sclerosis) there is enrichment in the CD3, CD4 and CD8 positive samples containing T cells as well as enrichment in the CD56 positive sample including NK cells. In addition, these disease SNPs overlap with hotspots present in the fetal thymus samples, consistent with the location of T cells maturation. Further signals specific to the individual aetiologies are also identified. Crohn’s disease SNPs show enrichment of overlap with hotspots in the fetal small and large intestine samples, as well as fibroblasts and skin cells. For inflammatory bowel disease SNPs there is a much more general enrichment, in addition to the specific immune cells, which may be consistent with the more generalized inflammation. In contrast, for autoimmune diseases where the primary involvement is a B cell response (Rheumatoid arthritis (RA), Systemic lupus erythematosus (SLE)), the most pronounced overlap enrichment is in CD19 positive samples characteristic of B cell activation or circulating plasma cells. In rheumatoid arthritis there is also some overlap enrichment for samples characteristic of T cells and thymocytes, but it is relatively less, and this is much less pronounced for SLE. Thus, FORGE analysis reveals tissue specific enrichment of overlap for GWAS SNPs with regulatory regions indicative of known tissue involvement in the disease aetiology.

**Figure 1.**
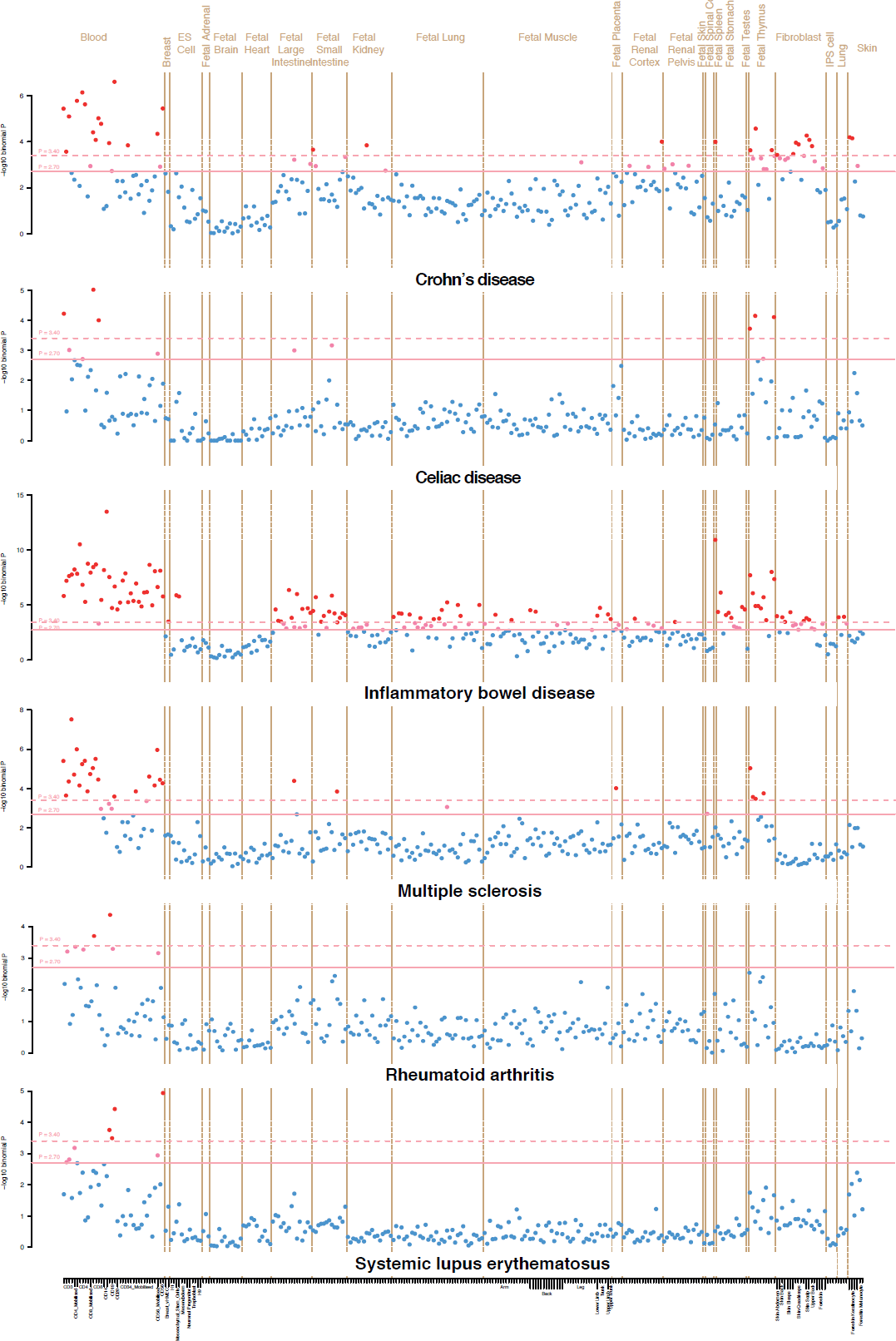
FORGE analysis results for GWAS of several autoimmune diseases on Roadmap Epigenome DNase I hotspots. A series of FORGE analysis results are presented for autoimmune phenotype GWAS SNPs. Each point represents the Z score (y axis) of the enrichment of the test SNP set compared to matched background SNPs on a single DNase I hotspot sample, organized by tissue as indicated by the brown labels at the top of the figure, and alphabetically by cell sample (x axis). Where informative, additional labels at the bottom of the figure highlight relevant distinct cell types. Red points are at Z >= 3.39 (empirical false positive rate <= 0.005 for 25 SNPs or more), pink points at Z >= 2.58. Full lists of the cells and results for each analysis are available in the Github data directory at https://github.com/iandunham/Forge. Phenotypes are labeled beneath each result.

The tissue specific enrichment of overlap is not specific to just autoimmune disease (see results gallery at http://www.1000genomes.org/forge-gwas-catalog-example-gallery11). For instance, for QRS duration the GWAS associated SNPs are strongly enriched for overlap with fetal heart samples. GWAS SNPs associated with pulmonary function measured by spirometry are enriched for overlap with hotspots in fetal lung cells and lung cell lines. For red blood cell traits and platelet count the major overlap enrichment signal is in CD34 positive hematopoietic progenitor cells consistent with their role in both red blood cell and platelet development. In contrast for GWAS SNPs involved in height, the overlap enrichment is not tissue specific but is more general over many tissues and cell lines. There are further examples displayed in the results gallery at http://www.1000genomes.org/forge-gwas-catalog-example-gallery11, in most case consistent with expected disease aetiologies.

## Discussion

FORGE (Functional element Overlap analysis of the Results of GWAS Experiments) analysis is a straightforward and fast method to examine sets of nucleotide variants, typically identified in GWAS studies, for tissue specific regulatory signals. It presents a graphical overview of overlap enrichment with DNase I hotspots that quickly provides evidence of a regulatory component to SNPs associated with a phenotype, and highlights potentially mechanistically relevant cells or tissues. A typical usage scenario would be to analyse a set of GWAS SNPs identified above genome wide significance to reveal the regulatory component of the association. Furthermore the cell or tissue enrichments might be consistent with prior expectation of the disease aetiology providing additional confidence in the SNP set identified, or might provide new insights as to potential sites of disease mechanism.

The statistical approach we used here relies on the careful matching of background behaviour of SNPs with calibrations by randomization for the per cell type enrichment. The underlying biases of GWAS SNP distributions with respect to TSS distance, maf and GC are not easy to model parametrically. However other approaches which would make assumptions of homogeneity (such as the Poisson distribution) or of regional heterogeneity (Genome Structure Correction, [31]) would not be able to capture these known biases. It is important to note that as alternate SNP resources are utilized in GWAS, the appropriate background SNPs must be used for control. For instance a switch to genotyping by genome sequencing or extensive use of imputation requires revision of the background. New and updated background sets can be implemented as required in particular for genome sequencing GWAS approaches and higher density genotyping arrays.

Not all GWAS study SNP sets downloaded from the GWAS catalog showed overlap enrichment with the DNase I hotspots. In these cases all sample points were above the P value thresholds (blue points). This could occur because there is no regulatory component underlying the GWAS association in these phenotypes, because the associated SNPs do not contain mechanistically causal SNPs, because the relevant tissue is not present in the available DNase I datasets or because of low power in the GWAS study to detect regulatory effects. As further data sets are release by the ENCODE, Roadmap Epigenome and other projects these can be incorporated into the database to provide coverage of further cell types. In addition the approach could be readily extended to other data types including regions of specific histone modification as used by Trynka et al [23] or relevant transcription factor binding regions.

60 out of 260 sets of GWAS SNPs from the GWAS catalog for specific phenotypes had overlap enrichments detected in at least one DNase I hotspot sample (Table 1). As described above in several cases, the patterns of tissue–specific enrichment are highly evocative of the known aetiologies of the phenotypes, but can also reveal additional tissue involvements that require further investigation. We encourage interested parties to peruse the gallery of results for their own phenotypes, as well as running new SNP sets discovered in GWAS either through the web interface or with the standalone software.

## Methods

### DNase I Data

ENCODE consortium hotspots [26] were obtained from ftp://ftp.ebi.ac.uk/pub/databases/ensembl/encode/integration_data_jan2011/byDataType/openchrom/jan2011/combined_hotspots/. Roadmap Epigenome DNAse1 sequencing tag alignments were obtained from http://www.genboree.org/EdaccData/Current-Release/experiment-sample/Chromatin_Accessibility/. The files used correspond to that part of the Gene Expression Omnibus (GEO) accession GSE18927 beyond the data use embargo date. These alignments were processed by the Hotspot (http://www.uwencode.org/proj/hotspot/) [28, 29] method with the default parameters to give hotspot and peak files. For this analysis we choose to use the hotspots which are regions of general DNase I sensitivity rather than peaks which are more similar to DNase I hypersensitive sites because, although the method works with peaks, hotspots reveal more tissue specific signal (data not shown). Cell and tissue assignments for each of the data sets were made using the decodings available from the ENCODE Data Coordination Center tables (https://genome.ucsc.edu/encode/cellTypes.html) or from the BioSamples database (http://www.ebi.ac.uk/biosamples/) sampleGroup SAMEG31306. A list of samples used is provided in Supplementary Table S1.

### GWAS SNP data

The complete collection of SNPs discovered in GWAS studies curated in the NHGRI GWAS Catalog [5] were downloaded from http://www.genome.gov/gwastudies/ [30](Accessed 3rd September 2014). SNPs were grouped according to the annotation provided in the Disease/Trait field and only sets with 5 or more non-redundant SNPs were retained. See Supplementary Table S2 for list of SNP sets analysed. A set of files of the SNPs included in analysis for each phenotype is available in the data directory of the GitHub release, https://github.com/iandunham/Forge. For Forge analysis SNP sets were further filtered by LD pruning removing all but one SNP from a set of SNPs at r^2^ >= 0.8 in the 1000 genomes data (see below) and were analysed for both ENCODE and Roadmap Epigenome DNase I hotspots, selecting background SNP sets from the default GWAS genotyping array SNPs.

### Preparation of FORGE overlaps

The FORGE tool utilises either an SQLite (command line tool, http://www.sqlite.org) or MySQL (web tool, http://www.mysql.com) database of the overlaps of every 1000 genomes project (http://www.1000genomes.org) [32]SNP with the ENCODE and Epigenome Roadmap DNase1 hotspots. To prepare this database, SNPs from the 1000 genomes phase 1 integrated call data set were downloaded from ftp://ftp.1000genomes.ebi.ac.uk/vol1/ftp/phase1/analysis_results/integrated_c_all_sets and compared to indexed DNase I hotspots using tabix from the SAMtools package (http://samtools.sourceforge.net/tabix.shtml) [33] using a distributed approach on the EBI compute farm. The overlaps for each SNP were stored in a single large indexed table of SNP location and identifier with binary strings representing the presence (1) or absence (0) of overlap in each sample for each of the hotspot data sets (ENCODE or Roadmap).

### Background SNP parameters

To prepare sets of background SNPs matched to the test SNP set, FORGE matches SNPs based on GC, maf and TSS distance, and repeats the overlap analysis for each of 1000 background sets. Overall population mafs were obtained from the 1000 genomes project phase 1 integrated call data set at ftp://ftp.1000genomes.ebi.ac.uk/vol1/ftp/phase1/analysis_results/integrated_call_sets. To control for the processes involved in selecting SNPs for genotyping, only 1000 genomes phase 1 SNPs that had been included on one of the common genotyping platforms as described in Ensembl (http://www.ensembl.org/info/genome/variation/data_description.html#variation_sets) were considered further. This left either 1875813 SNPs across various platforms (Affy GeneChip 100K Array, Affy GeneChip 500K Array Affy SNP6, HumanCNV370-Quadv3, HumanHap300v2, HumanHap550v3.0, Illumina Cardio Metabo, Illumina Human1M-duoV3, Illumina Human660W-quad) or 2231212 SNPs across the Illumina HumanOmni2.5 array. TSS distance was determined for each remaining SNP relative to the TSS defined by the Gencode project [34, 35] given in aType/gencode/jan2011/Gencodev7_CAGE_TSS_clusters_June2011.gff.gz using Bedops closest-features [36]. GC was determined for a 100 bp window centred on the SNP at base 50. The SNPs were then sorted into 1000 bins partitioned by deciles for each of GC, maf and TSS. For each SNP in a test set, the corresponding bin is identified based on its GC, maf and TSS distance, and background selections are made from that bin.

### FORGE analysis

A set of SNPs can be presented to FORGE as a list of RSIDs or by genome location on human genome build GRCh37 in either VCF or Bed formats. If RSIDs are not provided in one of these formats, the genome coordinates are used to identify the RSID. SNPs not present in the 1000 genomes phase 1 integrated call data set are excluded from the analysis. With LD pruning selected a single SNP (the first in the file) is chosen from LD clusters within either r^2^ >= 0.8 or r^2^ >= 0.1 as specified. For each analyzable SNP in the test set, overlaps are retrieved from the FORGE database, and a count of total hotspot overlaps is recorded for each DNase I sample (cell) for the test SNP set. One hundred matching background SNP sets containing the same number of SNPs as the test SNP set are selected, matched for GC, maf and TSS distance by decile bins as described above. Overlaps for each of the SNPs in each of the background SNP sets are also retrieved from the database and an overlap count for each background set in each DNase I sample is recorded. For each test SNP set, the background probability of overlap is determined from the 1000 background set overlap counts and the probability of the observed test result under a binomial distribution is calculated. The P value thresholds of 0.05 and 0.01 are corrected for multiple testing by division by the number of tissue groupings tested, and the corrected threshold is used. The use of tissue as the unit for sample grouping is consistent with the groupings obtained by hierarchical clustering of samples by Dnase 1 data (results not shown). The corrected thresholds are therefore more stringent than established by the random trials.

### FORGE outputs

FORGE generates both tabular and graphic descriptions of the enrichment of overlap for the test SNPs for each DNase I hotspot sample. A tab-separated values (TSV) file is output including columns for the binomial P value, cell, tissue, filename of the sample hotspots, SNPs that contribute to the enrichment, and the GEO accession for each sample. This data is also provided as an interactive table produced using the Datatables (https://datatables.net/) plug-in for the jQuery Javascript library accessed through the rCharts package (http://ramnathv.github.io/rCharts/).

Each of the graphic outputs presents the −log_10_ binomial p by cell sample. A pdf graphic is generated using base R graphics (http://www.r-project.org). The interactive Javascript graphic is generated using the rCharts package (http://ramnathv.github.io/rCharts/) to interface with the dimple d3 libraries (http://dimplejs.org). In both cases cells are grouped alphabetically by tissue, and for the pdf alphabetically by cell. The interactive graphic stacks replicate samples at the same x coordinate. In each of the graphics the colouring of results by P value is consistent, blue (p > 0.05 equivalent after correction), pink (0.05 => p < 0.01), and red (p <= 0.01). The corrected P value threshold is given on the pdf output.

### False positive rates

To estimate false positive rates, 1000 sets of SNPs at each of a series of SNP counts between 5 and 300 SNPs were randomly chosen from the 1000 genome phase 1 integrated SNP set. FORGE analysis was run for each set across the ENCODE and Roadmap Epigenome data, and the number of tests with P values less than thresholds ranging from 0.05 to 0.001 were recorded. These represent the false positives from 1000 trials at each of 424 samples i.e. 424,000 tests, and were used to calculate false positive rates at each significance threshold.

### Hierarchical Clustering of DNase I samples

The hierarchical clustering solution was obtained using a multi-scale bootstrap resampling approach. We first computed a binary regulatory signature for each cell type classifying each DNase I site as active or inactive in each cell type sample. Hierarchically clustering of the binary regulatory signatures was by Euclidean distance with Ward's agglomerative method using the pvclust R/CRAN package with default values (http://cran.r-project.org/web/packages/pvclust/index.html). Finally, we identified clusters supported by the data at a bootstrap probability p value < 0.01.

### Access and Source Code

FORGE is available through a web interface at http://browser.1000genomes.org/Homo_sapiens/UserData/Forge. The source code for FORGE is available on GitHub at https://github.com/iandunham/Forge with the Forge.db sqlite database and background selection hash tables availableat ftp://ftp.1000genomes.ebi.ac.uk/vol1/ftp/technical/browser/forge_11. FORGE has been successfully been installed and run on Mac OSX 10.8.4 and Red Hat Linux.

## Links

Web tool http://browser.1000genomes.org/Homo_sapiens/UserData/Forge

Web documentation http://www.1000genomes.org/forge-analysis-11

Results Gallery http://www.1000genomes.org/forge-gwas-catalog-example-gallery11

Source Code and Database https://github.com/iandunham/Forge ftp://ftp.1000genomes.ebi.ac.uk/vol1/ftp/technical/browser/forge_11

## Authors’ contributions

ID and EB designed the analysis and wrote the paper. ID wrote software and ran analysis. SM conducted the clustering of cell types by DNase I regions. EK implemented the web interface. VI provided statistical advice and discussion.

## Acknowledgements

Many thanks to Jan Quell for initial implementation of the pdf graphic and to Ramnath Vaidyanathan and @timelyportfolio for assistance with rCharts. We thank Prof Ajay Shah and Marc-Philip Hitz for comments on the results of cardiac phenotypes, and Graham Ritchie, and Nicole Soranzo for discussions.

## Supplementary Tables

Supplementary Table S1

Supplementary List of DNase I hotspot samples included in FORGE analysis. The table lists details of the 125 ENCODE and 299 Roadmap Epigenome samples in tab separated value format (tsv). The fields are

File: File name

Lab: The data-generating lab. One of UW (University of Washington, John Stamatoyannopoulos lab), Duke:UNC:UTA (Duke University, Greg Crawford lab) or combined representing a merged dataset from both labs.

Experiment type: always DNase-seq in the current implementation.

Project: Either ENCODE or Roadmap

Cell: The cell type

Tissue: Tissue name derived as described above.

Datatype: Always hotspots in the current implementation.

Short name: A short sample name used for plotting.

Individual: Either the code for the individual sample as described in Biosamples or NA if not available

GEO accession: The GEO accession where found, or “Not found” if it could not be deconvoluted.

Supplementary Table S2

Supplementary List of phenotypes analysed, with non-redundant SNP counts and Pubmed identifiers for the studies involved. This table is also available in the data directory of the GitHub release, https://github.com/iandunham/Forge.

